# Bacterial influence on the maintenance of symbiotic yeast through *Drosophila* metamorphosis

**DOI:** 10.1101/2020.05.31.126185

**Authors:** Robin Guilhot, Antoine Rombaut, Anne Xuéreb, Kate Howell, Simon Fellous

## Abstract

Interactions between microbial symbionts of metazoan hosts are emerging as key features of symbiotic systems. Little is known about the role of such interactions on the maintenance of symbiosis through host’s life cycle. We studied the influence of symbiotic bacteria on the maintenance of symbiotic yeast through metamorphosis of the fly *Drosophila melanogaster*. To this end we mimicked the development of larvae in natural fruit. In absence of bacteria yeast was never found in young adults. However, yeast could maintain through metamorphosis when larvae were inoculated with symbiotic bacteria isolated from *D. melanogaster* faeces. Furthermore, an Enterobacteriaceae favoured yeast transstadial maintenance. Because yeast is a critical symbiont of *D. melanogaster* flies, bacterial influence on host-yeast association may have consequences for the evolution of insect-yeast-bacteria tripartite symbiosis and their cooperation.

**Summary statement:** Bacterial symbionts of *Drosophila* influence yeast maintenance through fly metamorphosis, a novel observation that may have consequences for the evolution of insect-yeast-bacteria interactions.

## Introduction

Metazoans often form associations with non-obligatory symbiotic microorganisms. Microbial symbiont can influence host phenotype, and hosts determine symbiont multiplication and dispersal (Ferrari and Vavre, 2011). The importance of interactions between microorganisms is relatively better understood in the context of parasitism than in the context of beneficial symbionts (Alizon et al., 2013; Tollenaere et al., 2016; Zélé et al., 2018). Beneficial microbial symbionts can nonetheless interact in a wide variety of manners (Comolli, 2014; Seth and Taga, 2014; Hassani et al., 2018) and affect both the dynamics of each microorganism and the phenotype of their host (Wargo and Hogan, 2006; Newell and Douglas, 2014; Callens et al., 2018; Gould et al., 2018; Sommer and Newell, 2019).

Fungi-bacteria interactions are well described microbial interactions due to their importance in human health, industry and domestic life (Kobayashi and Crouch, 2009; Jouhten et al., 2016; Carbonetto et al., 2018). In the wild, yeasts and other fungi associate with extracellular bacteria in a wide variety of habitats, including decaying plant materials where they interact with the larvae and adults of saprophagous insects such as *Drosophila* flies. Symbioses between *Drosophila* and either yeast and bacteria have however been largely studied separately. Each type of microorganism affects *Drosophila* physiology, nutrition, reproduction and behavior (Ryu et al., 2008; Anagnostou et al., 2010; Shin et al., 2011; Storelli et al., 2011; Becher et al., 2012; Broderick and Lemaitre, 2012; Wong et al., 2014) and may maintain through - a variable part of - *Drosophila* life cycle (Bakula, 1969; Starmer et al., 1988; Hoang et al., 2015; Pais et al., 2018). In natural fruit, *D. melanogaster* larval development can be impossible, or largely compromised, in absence of yeast (Becher et al., 2012), suggesting symbiosis with yeast may not be dispensable for larvae. A handful of studies considering both fungi and bacteria showed that direct interaction between yeast and bacteria can modulate fly behavior (Fischer et al., 2017) and that bacteria can affect fly attraction to yeast (Leitão-Gonçalves et al., 2017). There is to date little data on how interactions between microbial symbionts affect their transmission from the host or maintenance among host life stages, which would have wide consequences for the ecological dynamics of the microorganisms, and consequently, the evolution of their symbiosis with flies.

It is established that *Drosophila* flies contribute to bacteria and yeast dynamics through effects on local multiplication and dispersal (Gilbert, 1980; Ganter, 1988; Starmer et al., 1988; Chandler et al., 2012; Stamps et al., 2012; Buser et al., 2014). *Saccharomyces* yeast, for example, attracts adult flies with volatiles, these adults then acquire the microorganism and later inoculate it in fruit where larvae develop. An alternative mean of insect vectoring for symbiotic bacteria and yeast of larvae would rely on the transstadial maintenance, from the larval to the adult stage. It would enable the colonization of freshly emerged adults by larval symbionts and, provided they maintain and are shed later in life, their possible dispersal to new patches of resources. The maintenance of yeast or bacteria throughout the *Drosophila* life cycle have been investigated (Rohlfs and Hoffmeister, 2005; Pais et al., 2018), including maintenance from larvae to adults through metamorphosis (i.e. transstadial maintenance or transstadial transmission) (Bakula, 1969; Starmer et al., 1988; Ridley et al., 2012; Duneau and Lazzaro, 2018). To our knowledge, how interactions between microbial symbionts affect symbiont maintenance through *Drosophila* life stages remains however unknown. Here, we investigated the maintenance of a wild isolate of *Saccharomyces cerevisiae* yeast from larvae to young adults *Drosophila melanogaster*. We found that yeast presence in adults only occurred when larvae were associated with symbiotic bacteria and its frequency depended on the identity of these bacteria.

## Material and methods

### Biological material

We used a *Drosophila melanogaster* (Meigen, 1830) Oregon-R strain usually maintained on banana medium (233 g.L^-1^ banana, 62 g.L^-1^ sugar, 62 g.L^-1^ dead yeast, 25 g.L^-1^ ethanol, 10 g.L^-1^ agar and 5 g.L^-1^ nipagin) at 21°C with a 14 h/10 h day-night cycle. The four bacterial strains used had been isolated from feces of adult Oregon-R flies and had been described in Guilhot et al. (2019). Bacterial strains were identified as *Staphylococcus* sp. (accession number MK461976 in the NCBI database), *Enterococcus* sp. (MK461977), an Enterobacteriaceae (MK461978) and an Actinobacteria (MK461979). The taxonomic resolution of our analyses is unfortunately modest and these bacteria do not usually dominate the *Drosophila* microbiota. However similar strains had already been identified as associated with Drosophilids (Chandler et al., 2011; Staubach et al., 2013). More importantly, microbial species and strains can evolve rapidly and with large consequences on their effects on host phenotype (Winans et al., 2017; Martino et al., 2018). For this reason, the proper description of symbiont effects on host phenotype in relevant experimental conditions (Guilhot et al. 2019) may be more important than high taxonomical resolution to understand symbiosis.

The yeast *Saccharomyces cerevisiae* (Meyen ex Hansen, 1883) strain was isolated from a wild Drosophilid in the *‘Le Domaine de l’Hortus’* vineyard, near Montpellier in Southern France.

### Experimental design

The main experiment was conducted on sterile vials. Each experimental unit consisted of twenty *D. melanogaster* eggs free of cultivable bacteria and yeast deposited on an artificial wound of a surface-sterilized grape berry that was inoculated, or not, with specific microorganisms. Eggs were laid by conventionally-reared *D. melanogaster* Oregon-R females on solidified grape juice, which contained the antibiotic streptomycin (1 mg.L^-1^, from a standard streptomycin solution of 1 mg.ml^-1^ in 1 mM EDTA (Sigma-Aldrich ref. 85886)) in order to remove parental bacteria. Grape berries were surface-sterilized; they were hence dipped in a 2% bleach solution, rinsed with sterile water and dried before use. We ensure these procedures were efficient: no cultivable bacteria or yeast were found in surface-sterilized berries and eggs homogenates. Every grape berry received 10^4^ *S. cerevisiae* yeast cells suspended in 10 μl. Berries inoculated with *S. cerevisiae* were then allocated to six different bacterial treatments: no bacteria (control, n = 18); one of the four bacterial strains described above (10^4^ cells of the same type: n (*Staphylococcus*) = 13, n (*Enterococcus*) = 13, n (Enterobacteriaceae) =11, n (Actinobacteria) = 13); and a mixture of the four bacteria (2.5×10^3^ cells of each type, n = 9). Replicates were organized in eleven blocks launched over four days.

Newly formed pupae were aseptically removed daily from their larval container and placed in a sterile new vial until adult emergence. This procedure mimicked natural insect behavior as *D. melanogaster* larvae usually crawl out of their substrate before pupation (Sokolowski et al., 1986; Woltz and Lee, 2017), which incidentally prevent the exposure of young adults to the microorganisms present in the larval substrate. It was not logistically feasible to process every adult independently. We therefore elected to assess microbial content in groups of adults that emerged on the same day. For each grape berry, we randomly selected a single pupa and pooled all the adults, females as males, that emerged on the day as this pupa. This protocol allowed the detection of yeast (and bacterial) cells associated with freshly emerged adults that may have been present externally or internally. Sampled adults were homogenized in sterile PBS using a Tissue Lyser II (Qiagen) and Ø3 mm glass balls. Serially diluted fly samples were plated on Lysogeny Broth (LB) plates to count microbial cells after incubation at 24°C. Colonies of the five microbial symbionts (yeast and bacteria) were distinguished according to their morphology (see Guilhot et al. 2019).

Effects of bacteria on yeast development in the larval fruit may explain some of our results. We hence collected the remaining juice from grape berries two days after the formation of the last pupa. Although quantifying yeast in fruit when larvae were feeding would have been informative too, we elected to not disturb the development of the larvae and waited the larvae left their substrate. Serially diluted fruit samples were plated on LB plates to count yeast colonies at 24°C. As above, microbial colonies were distinguished based on their morphology.

In parallel to the main experiment, bacteria were also inoculated without yeast on cubes of laboratory medium (which contains dead yeast) following the procedure above to assess bacteria transstadial maintenance in their environment of origin (i.e. the banana medium used to rear the fly colony and described above). The six different bacterial treatments were: no bacteria (control, n = 12); one of the four bacterial strains described above (n (*Staphylococcus*) = 12, n (*Enterococcus*) = 7, n (Enterobacteriaceae) = 11, n (Actinobacteria) = 11); and a mixture of the four bacteria (n = 14). Replicates were organized in fifteen blocks launched over four days.

### Statistical analyses

We hypothesized larval bacteria would affect yeast transstadial maintenance. We estimated this phenomenon in groups of 1 to 11 freshly emerged adults (median = 5, IQR = 4). Yeast-positive samples contained 1 to 150 cells per adult fly (Fig. S1). This variation was not investigated statistically due to low statistical power. Whether live yeast cells were present or not was analyzed using a generalized linear model with binomial distribution and logit link function. Tested factors comprised bacterial treatment, number of adults in the groups, yeast concentration in the fruit, age of the adults and experimental block. Statistical power did not enable testing the interaction between bacterial treatment and number of adults in the groups (but see Fig. S2). Because we used groups of adult flies it was mandatory to take into account the number of individuals per pool. The biological material employed (i.e. wild and laboratory strains and populations) informs on the factors that can influence transstadial symbiont maintenance in a qualitative fashion, but could not indicate their quantitative occurrence in the field. Backward model selection allowed to eliminate non-significant terms (i.e. yeast concentration in the fruit and age of the pooled flies) from the initial complete model. Contrasts were used to detect significant differences between bacterial treatment levels. Numbers of replicates varied among bacterial treatments due to differential larval mortality. However, the analysis of larval survival revealed no significant effect of the bacteria on this trait (Guilhot et al., 2019).

To study the effect of the bacterial treatment on the yeast concentration (log-transformed) in larval fruit substrate, we used a linear mixed model with Restricted Maximum Estimate Likelihood. Experimental block was defined as a random factor.

Analyzes were performed with JMP (SAS, 14.1).

## Results

Bacterial treatment significantly affected *S. cerevisiae* presence in freshly emerged adult flies (χ^2^ = 20.30, df = 5, p = 0.001). Yeast was never observed in adult flies that emerged from control treatments, unlike treatments with bacteria at the larval stage (contrast ‘All treatments with bacteria’ vs ‘Control’: χ^2^ = 11.2, df = 1, p < 0.0001) (Fig. 1). Young adult flies that had developed with the Enterobacteriaceae alone or in mixture with the other bacteria were more likely to harbor live yeast cells than the other treatments with bacteria at the larval stage (contrast ‘With Enterobacteriaceae’ vs ‘All other treatments with bacteria’: χ^2^ = 4.52, df = 1, p = 0.03) (Fig. 1). As expected, the number of individuals in the assayed pool significantly and positively affected the likelihood of yeast observation (χ^2^ = 7.54, df = 1, p = 0.01) (Fig. S2) – supporting the need to include this factor in all the analyses. The age of freshly emerged adult flies (χ^2^ = 0.65, df = 1, p = 0.42) and the yeast concentration in the larval medium (χ^2^ = 0, df = 1, p = 1) had not significant influence on yeast presence in adults.

**Fig. 1.**
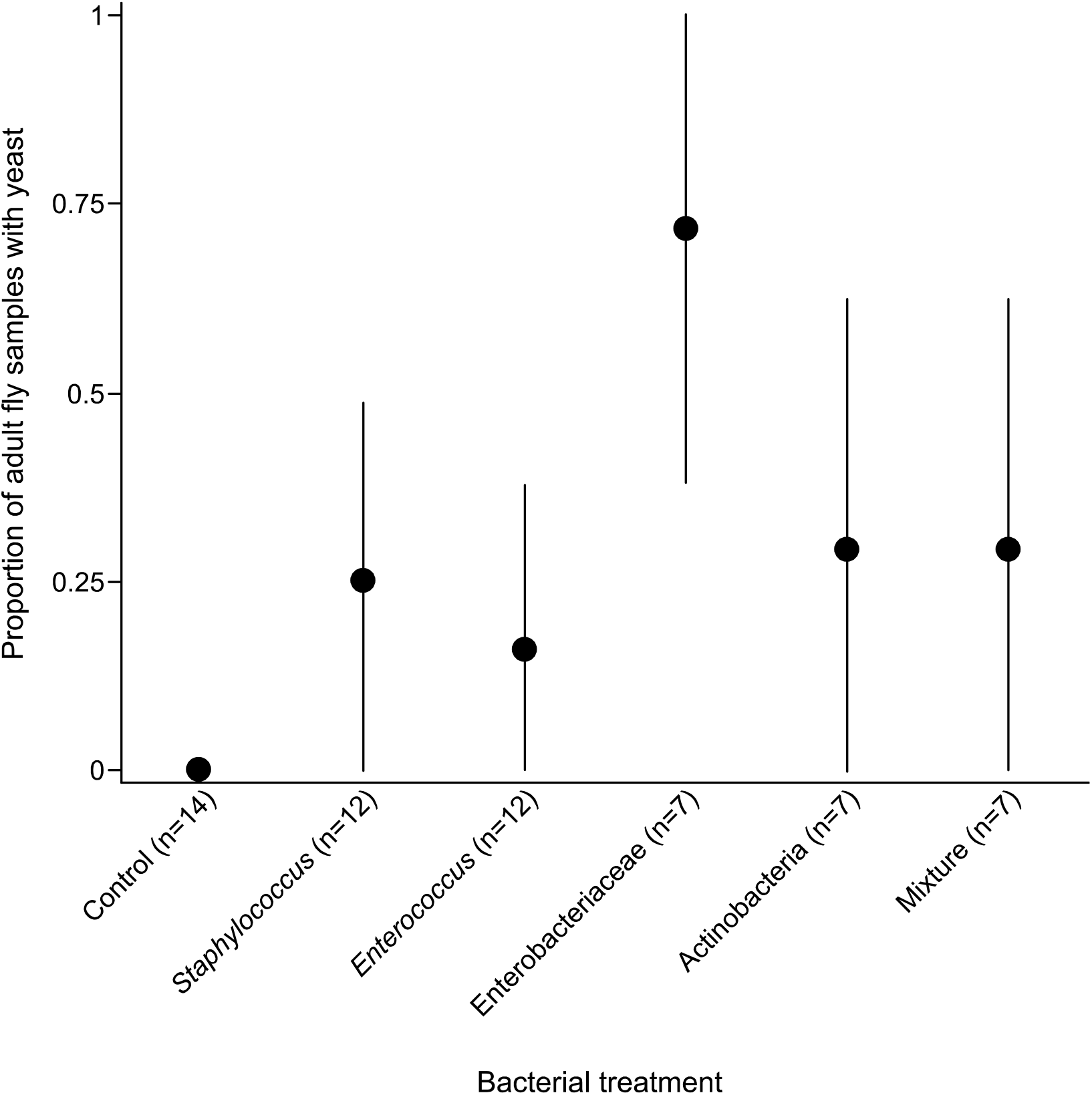
Transstadial maintenance of *Saccharomyces cerevisiae* in response to bacterial treatment. Symbols indicate the proportion of groups of freshly emerged adult flies containing yeast per bacterial treatment (n = number of adult groups per bacterial treatment). 95% binomial confidence intervals were calculated using normal approximation method. These results are qualitative as we used groups of adult flies to estimate yeast transstadial maintenance (Fig S2).

The bacterial treatment had no significant effect on the yeast concentration in the medium two days after the formation of the last pupa (F_5,49_ = 1.18, p = 0.33) (Fig. 2). Yeast presence in fruit flesh was detected in all replicates but one.

**Fig. 2.**
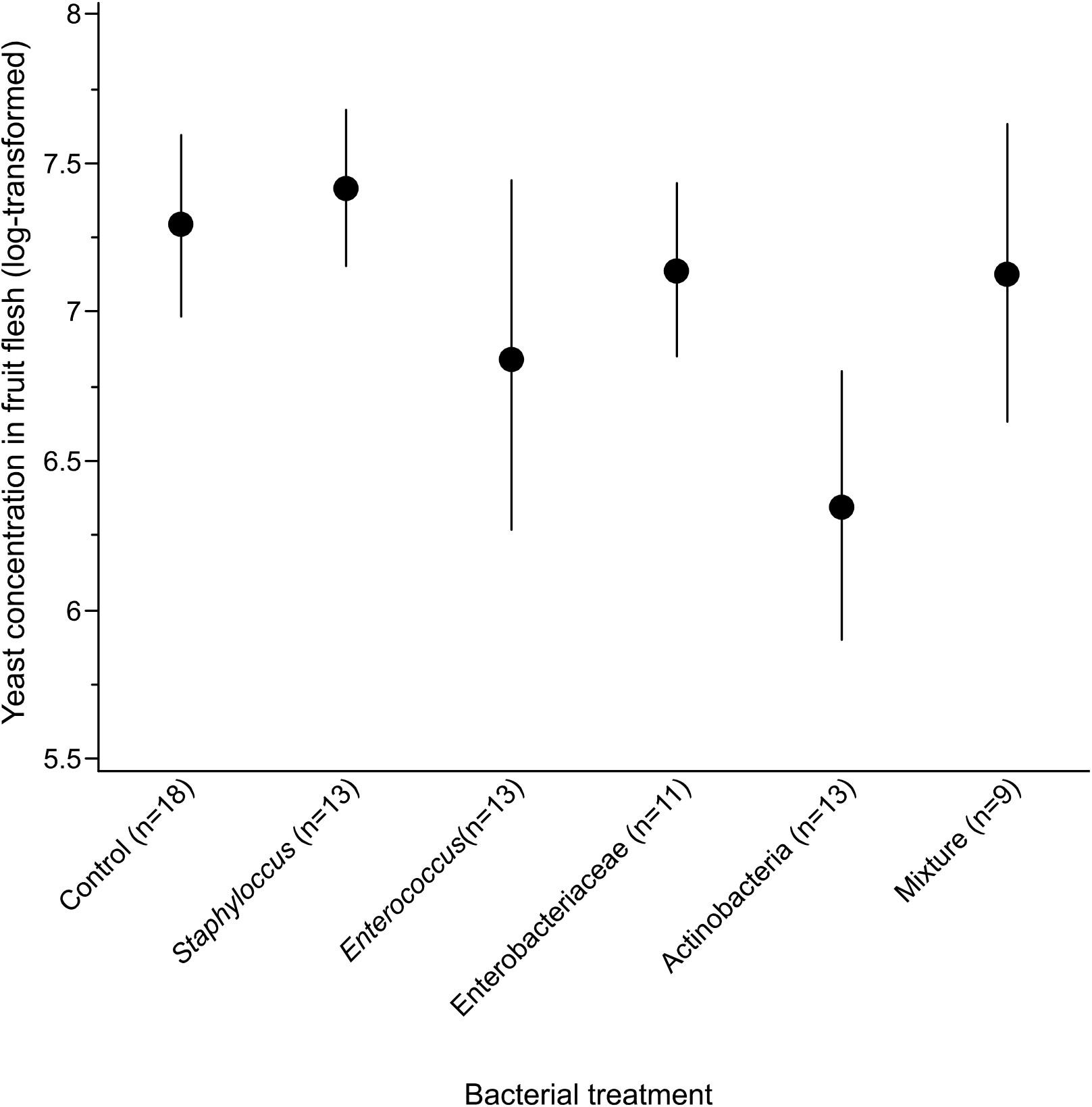
Yeast concentration in grape berry flesh after the formation of the last pupa. Concentration is expressed in number of yeast cells per 200 μl of fruit flesh. Symbols indicate mean ± s.e.m.

Bacteria could be observed in young adults that emerged from most combinations of larval environment and bacterial treatment (Fig. S3). However, the proportion of bacteria-positive groups never exceeded 25%. When bacteria were detected, load varied from 1 to 33 bacterial cells per adult fly (data not analyzed statistically). The observation of the inoculated bacteria in emerged adults shows these bacteria sampled in laboratory adults reared on artificial medium could establish symbiosis with larvae, even in fruit substrate.

## Discussion

Our most important result is that larval bacteria influenced yeast transstadial maintenance (Fig. 1). In control treatments that were not inoculated by bacteria, yeast was never found in freshly emerged adult flies. On the contrary, the presence of bacteria at the larval stage favored yeast maintenance through host metamorphosis. In particular, inoculation by the Enterobacteriaceae bacterium (alone or in mixture) led to greater *S. cerevisiae* transstadial maintenance than the other bacterial treatments (Fig. 1). The propensity to favor yeast maintenance hence seemed to vary among bacteria.

It is well known that coinfecting symbionts (mutualistic as parasitic) often affect each other’s horizontal transmission to new hosts in holometabolous insects (Azambuja et al., 2005; Fellous and Koella, 2009; Gendrin and Christophides, 2013; Hegde et al., 2015) and other multicellular organisms (Azambuja et al., 2005; Lass et al., 2013; Barret et al., 2016; Bonnet et al., 2017; Zélé et al., 2018). However, we know a single other case of microbial interactions affecting symbiont maintenance through complete metamorphosis: in *Galleria mellonella* butterflies, the bacterium *Enterococcus mundtii* interacts with host immunity during the pupal stage to shape adult bacterial microbiota (Johnston and Rolff, 2015). Although we used fresh fruit and a wild yeast strain, flies and bacteria were laboratory sourced. Our experiment hence shows bacteria affect yeast transstadial maintenance in *D. melanogaster*, but further work will be necessary to unveil the pervasiveness of this phenomenon in the field.

What mechanisms may underlie symbiont transstadial maintenance, and how did bacteria affect it? The maintenance of *S. cerevisiae* yeast and several bacterial strains through *Drosophila* metamorphosis are congruent with previous reports of the transstadial maintenance of extracellular microbial symbionts in Drosophilids (Bakula, 1969; Starmer et al., 1988; Ridley et al., 2012; Duneau and Lazzaro, 2018; Téfit et al., 2018), other Dipterans (Radvan, 1960; Capuzzo et al., 2005; Rochon et al., 2005; Damiani et al., 2008; Lauzon et al., 2009; Gendrin and Christophides, 2013; Nayduch and Burrus, 2017; Majumder et al., 2020) and other holometabolous insects (Hammer and Moran, 2019). Microbial symbionts could maintain on inner or outer walls of the pupal chamber (Kaltenpoth et al., 2010; Wang and Rozen, 2017). In *Drosophila melanogaster*, bacterial cells of *Escherichia coli* were found associated with the inner pupal membrane (Bakula, 1969). Alternatively, adults might retrieve symbionts by consuming their own meconium - the remaining of larval midgut that is excreted right after adult emergence (Moll et al., 2001; Broderick and Lemaitre, 2012; Gendrin and Christophides, 2013). The mechanism of bacterial influence on yeast maintenance through metamorphosis is not trivial either. The Enterobacteriaceae that favored yeast maintenance, despite presenting a wide metabolic spectrum (Guilhot et al., 2019), probably not improved fruit quality by concentrating or synthetizing nutrients (Ramiro et al., 2016) as had no significant effect on fly phenotype in this context (Guilhot et al., 2019). Besides, the concentration of yeast cells in fruit did not correlated with the presence of yeast in the freshly emerged adults and was not affected by the bacterial treatment (Fig. 2). This lack of quantitative relationships suggests yeast maintenance through metamorphosis may be determined by qualitative processes rather than mere cell numbers. Several bacteria are known to interact with *Drosophila* host signaling (e.g. Shin et al., 2011; Storelli et al., 2011). Symbiotic bacteria could therefore elicit host or yeast physiological responses in a way that would affect the likelihood of transstadial maintenance.

Yeast transstadial maintenance in *D. melanogaster* may have consequences for the spatial spread of the yeast and the evolution of the symbiosis. Yeast needs active transport by insect vectors to disperse among the ephemeral patch of resources formed by fruits (Starmer and Lachance, 2011). *Drosophila* adults contribute to such yeast dispersal through two non-excluding mechanisms. First, it is well established that yeasts produce chemical volatiles that attract adult flies (Palanca et al., 2013; Buser et al., 2014; Scheidler et al., 2015; Anagnostou et al., 2016; Bellutti et al., 2018; Günther et al., 2019; Lewis and Hamby, 2019), which favors their acquisition and vectoring by insects to new resource patches (Buser et al., 2014). Whether this phenomenon reflects yeast adaptation to insect vectoring is however debated (Günther and Goddard, 2019). Second, yeast maintenance through *Drosophila* metamorphosis - as demonstrated here - would enable the dispersal to new resource patches of larval symbionts (e.g. fruit, possibly infested with insect larvae) by colonized emerging adults. Such continuity in symbiosis over the life-cycle selects larval symbionts for beneficial effects on host fitness (Ebert, 2013). The microbial strains the most beneficial to larval development (for example in terms of larval survival) would be the ones best dispersed to new resources patches by the vigorous or numerous adult hosts they favored the development of. Furthermore, the maintenance of larval microbial symbionts until adult emergence may also benefit the host as freshly emerged adults could be less susceptible to opportunistic pathogens due to symbiont prior presence (Blum et al., 2013; Johnston and Rolff, 2015; Obadia et al., 2017). To conclude, transstadial maintenance of larval symbionts has implication for the dynamics and evolution of both hosts and microorganisms, the effects of bacteria on yeast we report may therefore affect important aspects of symbiosis. These are new and anticipated consequences of insect association with bacteria.

Symbiont-symbiont interactions are emerging as key features of numerous taxa (Ferrari and Vavre, 2011; Álvarez-Pérez et al., 2019, Mathé-Hubert et al., 2019), including *Drosophila* flies (Fischer et al., 2017; Gould et al., 2018). Studying microbial symbionts one by one may be more tractable, however our experiment illustrates understanding the nature and diversity of host-symbiont relationships necessitates encompassing the complexity of natural communities.

## Acknowledgements

We thank Edouard Jurkevitch, Natacha Kremer and Elodie Vercken for critical reading of an earlier version of this manuscript and Laure Benoit, Marie-Pierre Chapuis, Romain Gallet and Philippe Gautier for methodological help.

## Author contributions

R.G., A.R. and S.F. designed the experiment. R.G., A.R., A.X. and S.F. ran the experiment. K.H. collected the yeast isolate. R.G. and S.F. analyzed the data and wrote the manuscript.

## Data availability

The dataset is available in the open data repository Zenodo (doi: 10.5281/zenodo.3546129) (Guilhot et al., 2020).

