## Supplementary figures for "Bacterial influence on the maintenance of symbiotic yeast through *Drosophila* metamorphosis"

### Supplementary Material

**Fig. S1. Number of yeast cells per freshly emerged adult fly (log-transformed).**

Filled dots indicate mean ± s.e.m and open dots indicate individual values.

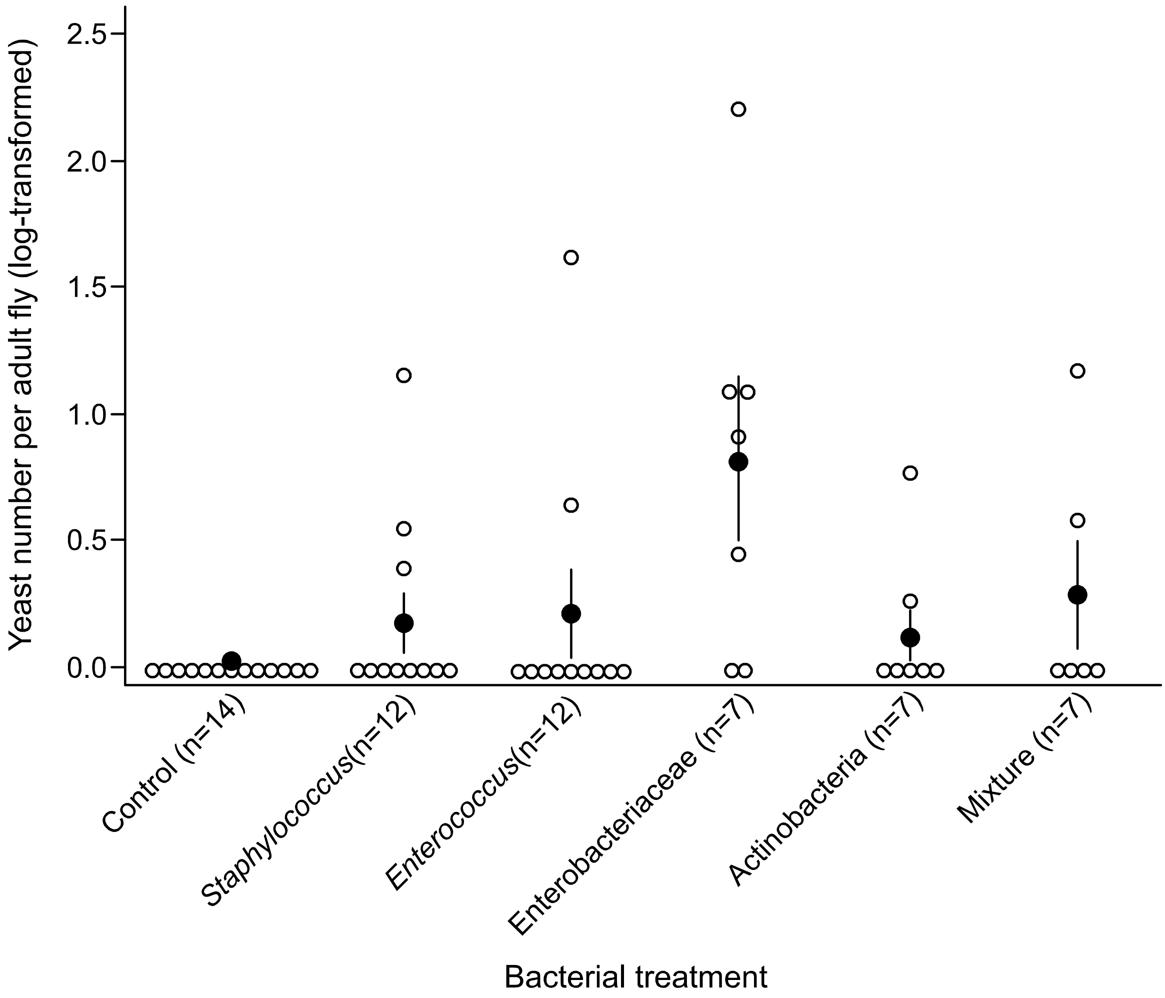

**Fig. S2. Relationship between the number of young adult flies in the groups and the likelihood of yeast transstadial maintenance.** The number of freshly emerged adult flies in the groups significantly and positively affected yeast detection ($\chi^2 = 7.54$, df = 1, p = 0.01). Filled dots indicate the proportion of adult groups containing yeast per number of adult flies in the groups (all treatments together). 95% binomial confidence intervals were calculated using normal approximation method. Other symbols indicate the proportion of adult groups containing yeast per number of adult flies in the groups for each bacterial treatment.

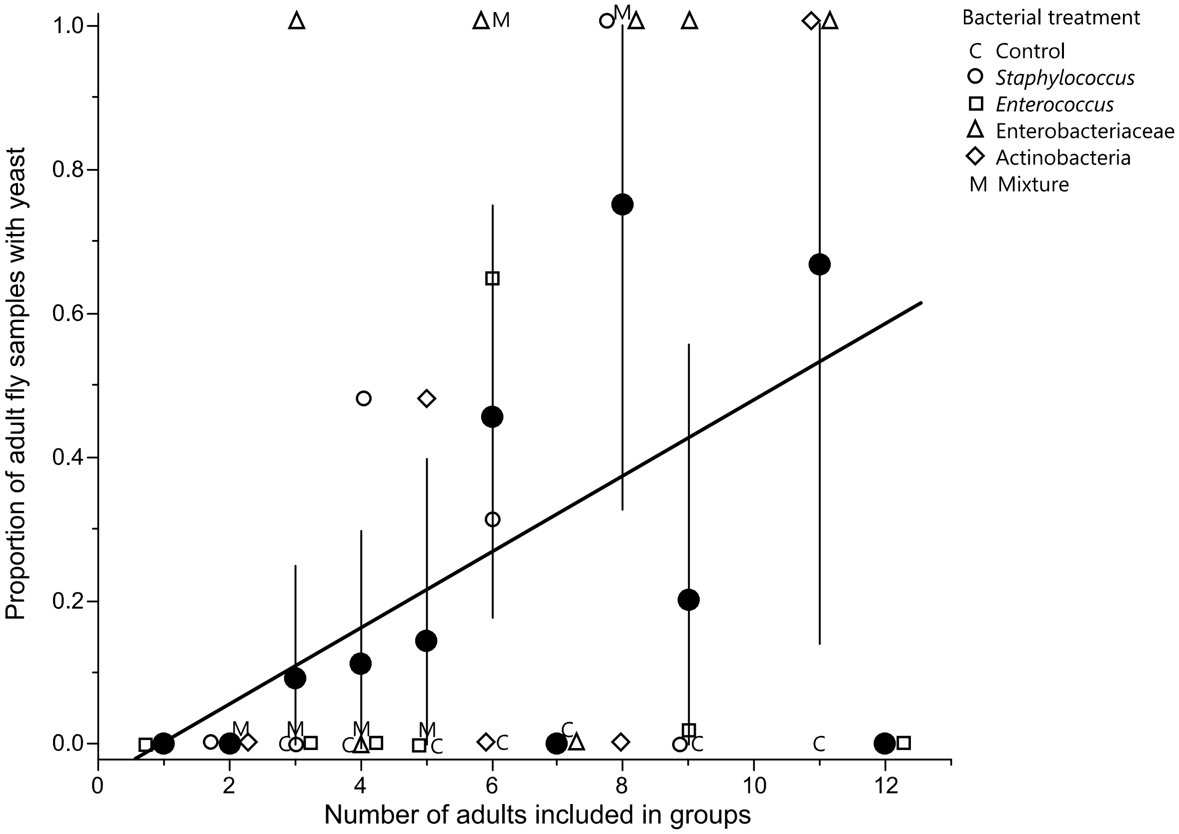

**Fig. S3. Transstadial maintenance of bacteria in grape berries (A) and in laboratory medium (B).** Symbols indicate the proportion of pools of freshly emerged adults containing bacteria for each bacterial treatment (n = number of pools). 95% binomial confidence intervals were calculated using normal approximation method.

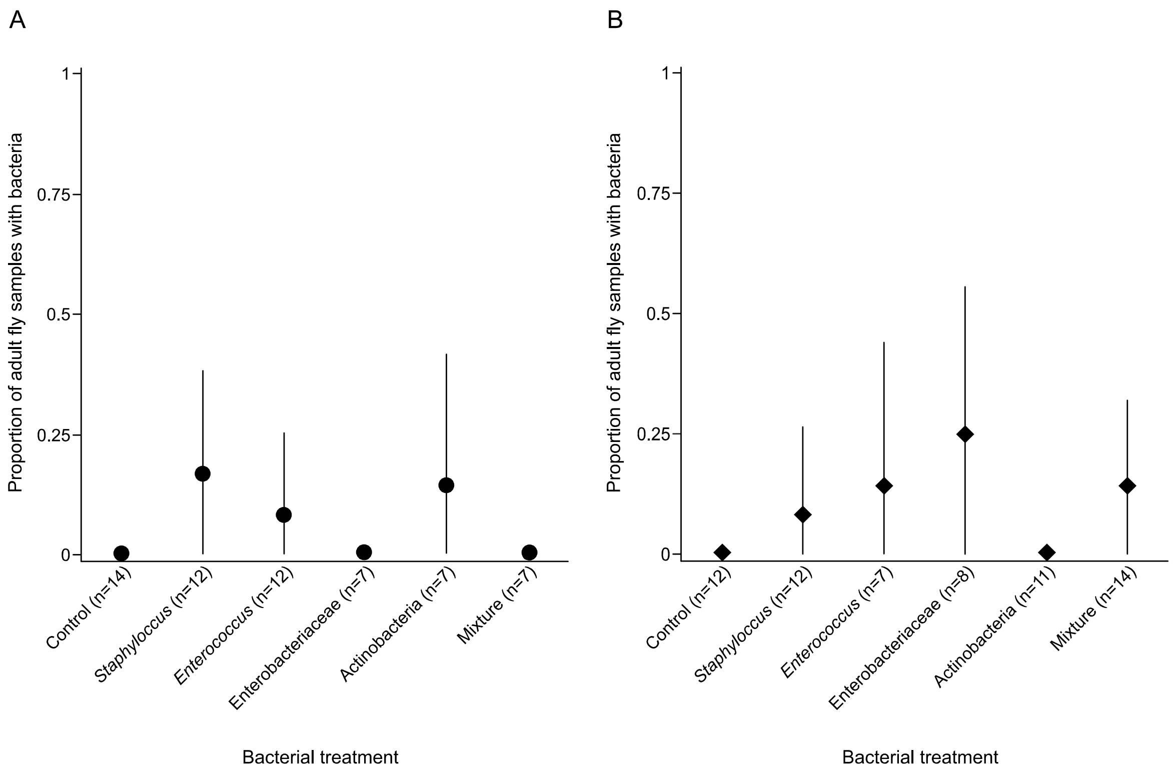
